# Testing for trait lability at macroevolutionary scales: is life history evolution labile in grasses?

**DOI:** 10.64898/2026.02.13.705712

**Authors:** Laura Schat, Aelys M. Humphreys

**Author notes:** Authors for correspondence: Laura Schat and Aelys M. Humphreys.

## Abstract

Trait lability has been defined in a myriad of ways in the macroevolutionary literature, yet there is no clear consensus what exactly is meant by a labile trait or how best to test for it. Here, we compare four frequently used approaches, using life history evolution in grasses as an example: i) phylogenetic signal, ii) ancestral state reconstruction with the hidden rates model, iii) ancestral state reconstruction with the threshold model and iv) simulations. We asked whether inferences of trait lability are consistent among approaches, and whether life history continues a labile trait in grasses. We found that inferences varied across the four approaches, with no two methods consistently yielding the same result. We advocate the use of simulations for comparing observed transition rates to expectations under stable and labile conditions. We infer numerous origins and losses of annuality in grasses, with annuals being unevenly distributed among clades (0-30%), concentrated in (sub)tropical clades comprising C_3_ and C_4_ species. Surprisingly, life history is not labile throughout grasses, but stable in the C_3_, temperate Pooideae. Our findings highlight the importance of objective tests of trait lability, based on predefined criteria, with implications for the field of macroevolution and beyond (e.g. crop breeding).

## Introduction

Every ecological, morphological and physiological trait has its own unique, evolutionary history. Although no two traits have evolved in exactly the same way, there are general patterns in how traits evolve. Some traits are labile and undergo a lot of evolutionary change (e.g., traits that evolve numerous times, or are gained and lost repeatedly), while other traits are stable, undergoing limited evolutionary change (e.g., traits with one or few origins with little subsequent change). Understanding how traits change, and whether a trait is labile or stable, can provide insights into the evolutionary processes that drive trait and organismal diversification. Specifically, quantifying trait lability can provide three key insights. Firstly, estimates of lability can contribute new insight into the predominant mode of evolution, i.e. whether evolution predominantly occurs gradually or in short bursts [1,2]. Secondly, the phylogenetic position of the origin of a labile trait can help determine the causes of evolutionary lability [3,4]. This underlying cause may be in the form of a precursor trait, facilitating multiple changes in the state or lability of another trait [3–6]. Thirdly, variation in the lability of a trait within a species or clade can be used to predict how they will perform under certain conditions. For example, trait lability indicates evolutionary flexibility, which can be incorporated into estimates of the impacts of climate change and anthropogenic pressure [5,7], or indicate which crops may be able to establish sought-after traits, such as perenniality and salt tolerance [5].

But what exactly is meant by trait lability? Although there are many methods available that directly or indirectly test lability of discrete traits, there are no clear criteria for defining what constitutes a labile trait, or how best to test for it. In some studies, lability is defined as a plastic trait that may change within the lifetime of an individual [8,9]. In others, lability refers to a traits which are constant within individuals, but vary within species [7]. Here, we focus on trait lability at macroevolutionary scales, i.e., on traits that may change (frequently) over at least several generations, and among closely related species, but that are assumed to be constant within individuals and species. Lability has been defined in a myriad of ways in the macroevolutionary literature, ranging from quantifiable statistics to more intuitive interpretations. One straightforward way of comparing the lability of multiple traits is by counting the number of state changes in ancestral state reconstructions (e.g. gains and losses for a binary trait; e.g. [5,10–12]).

According to this approach, salt tolerance is considered labile in grasses because it has originated 76 times and has few taxa per origin [5]. Similarly, C4 photosynthesis and bilaterally symmetrical flowers are considered labile in flowering plants as there have been more than 60 or 130 origins, respectively [13,14]. In some cases, there is no specific number of state changes associated with statements of lability. For example, flower shape in *Jaborosa* (Solanaceae) is considered labile due to its “repeated and independent changes”, growth form in *Crassula* is considered labile as it was “estimated to evolve multiple times”, and fang traits are considered labile in snakes because they underwent “multiple losses” [15–17]. This suggests that an implicit interpretation of lability is any trait that has been gained or lost more than once. Thus, the literature abounds with examples of evolutionary lability being used to describe traits that transition frequently between different states over the course of multiple generations or over branches in a phylogenetic tree [11,18,19].

However, inferring lability based on an absolute or even unspecified number of observed state changes can be ambiguous, as this number can vary with the size, age and sampling of a clade, and results may be sensitive to how a trait is scored or measured [20]. What is a reasonable cut-off for distinguishing labile from stable traits? What is a reasonable null expectation for trait lability? Comparing the number of state transitions among traits or clades, or with an evolutionary null scenario, can put the (relative) lability into context. For example, with over 70 origins in grasses, salt tolerance is more labile than C4 photosynthesis, with over 20 origins [5,21]. In *Solanum*, a variety of morphological traits have been ranked based on their mean number of observed state transitions, from those that are ‘highly evolutionary labile’ ( >100 transitions), through ‘evolutionarily labile’ (50-100 transitions) and ‘conserved’ (10–49 transitions) to ‘highly conserved’ (< 10 transitions) [12]. Similarly, growth form in Montiaceae has been referred to as labile, as there have been more transitions in this clade than in its sister clade [11]. Finally, salt tolerance is considered labile in over 20 angiosperm families, as there have been more frequent origins than expected under a null model (Brownian motion, BM; [10]). These examples allow inference of trait lability in particular comparative contexts, but do not necessarily allow general conclusions about the causes and correlates of trait lability beyond these contexts.

Evolutionary transition rates estimated under a given evolutionary model have also been used to infer lability, with higher rates leading to more frequent state changes, thus indicating lability. One class of models that may be particularly suited to inferences of lability are hidden Markov models (HMMs; [4]), because they allow evolutionary transition rates to vary in different parts of a phylogenetic tree. Inspired by models of nucleotide sequence evolution with site-specific rates (e.g. [22]), these models allow inference of how the (relative) lability of a trait varies among clades and over time. For example, growth form evolution in Montiaceae has been found to be both fast and slow, compared to just slow in their sister clade (a broad clade comprising several families including Cactaceae), thus contributing to the conclusion that growth form is labile in Montiaceae [11]. Similarly, this method has been used to show how growth form evolution varies across campanulids, being stable in Aquifoliales (low transition rates) and labile in Asteraceae (varying transition rates) [4]. Finally, this model has been used to distinguish regular and stable nitrogen fixing legumes [23]. Stable nitrogen fixers were defined as clades that are extremely unlikely to lose the nitrogen-fixing symbiosis, with the rate of loss being almost 60 times higher in the regular than the stable nitrogen fixers.

Another approach for estimating trait lability is the threshold model. With origins in quantitative genetics, it has been extended for comparative purposes to model discrete character evolution as a function of an underlying (unseen) liability factor, which itself evolves according to a model conditioned on the tip states (observed) of the discrete trait [24–26]. The state of the discrete trait is determined by the value of this liability factor: when the liability factor crosses a certain threshold, the discrete trait’s state changes. The closer the liability factor is to the threshold, the more likely it is to cross it, i.e. the more likely the discrete trait is to undergo a state change. Thus, the liability factor measures a discrete trait’s liability to change. When the liability factor is far from the threshold, a larger shift in value is required to cross the threshold and the discrete trait is less liable to change. This might be expected for a species or clade that has existed in a given state for a long time, due, perhaps, to other morphological, physiological or anatomical features associated with the current state [26]. The method has been widely adopted for estimating ancestral states of discrete traits (e.g. [27]), but the liability factor also allows inference of areas of a phylogenetic tree that are more or less prone to change, i.e., that are labile or stable. For example, life history strategy in Montiaceae was found to have liability factors closer to the threshold compared to its sister clade, i.e. to be more labile [11].

Lastly, estimates of phylogenetic signal have been used as proxies for trait lability. Phylogenetic signal is the tendency of closely related species to resemble each other more than a set of species sampled randomly from the same phylogenetic tree. High phylogenetic signal is often interpreted as trait stability and low phylogenetic signal as lability. However, it can be hard to determine lability based on a single estimate, so these are often compared to a distribution of values simulated under null scenarios, such as BM (i.e., high phylogenetic signal indicative of trait stability) or randomness (i.e., low phylogenetic signal indicative of trait lability). Such comparisons can distinguish whether estimates of phylogenetic signal indicate trait stability, lability or something in between. For example, this approach was used to test several drought related traits in *Salix* [28]. Three traits had higher phylogenetic signal than expected under a randomized null and are therefore more stable than expected by chance. Two traits had lower phylogenetic signal than expected under BM and are therefore more labile than expected under this model.

From the above is clear that trait lability has been inferred at macroevolutionary scales using a range of approaches and that most inferences are relative, e.g., to other traits, clades or expectations under a given null scenario. However, it remains unclear how consistent inferences of lability are among approaches and, as a consequence, whether previous inferences bear significance in a broader context, or are specific to the group and analytical approach studied. This is important in an evolutionary context, but also beyond, if we are to incorporate estimates of evolutionary flexibility into climate change forecasts or development of future crop varieties. Here, we compare inferences of trait lability across several approaches, using life history evolution in grasses (Poaceae) as a case study. Life history strategy (annual/perennial) is frequently referred to in the literature as one of the most labile traits of plants [19,29–31], generally being more labile in plants than in animals [32]. Annuals comprise only around 6% of vascular plant and 10% of angiosperm species [19,30,33]. However, this small proportion of species is not restricted to a small proportion of the lineages; annuals occur in all major clades of flowering plants [30]. Thus, angiosperms are thought to have transitioned from their ancestral perennial state to being annual frequently [6,11,30,34–38], a process that could have occurred “thousands of times” [19].

One angiosperm family with more annual species than almost any other are the grasses (Poaceae; [30]). Amongst these annuals are some of the world’s major food crops, such as wheat (*Triticum aestivum*), corn (*Zea mays*) and rice (*Oryza sativa*). To date, however, the origin and evolution of the annual life history strategy has not been modelled across the family as a whole, which would provide a broader evolutionary perspective and allow comparison of the timing and mode of evolution among clades. It is thought that the ancestor of Poaceae was a herbaceous perennial, as were the ancestors of the largest subfamilies, Pooideae and Panicoideae, with multiple transitions to annualism having been reconstructed in some clades (e.g. Danthonioideae and Pooideae; [6,38,39]). Within the Pooideae alone, there have been an estimated (at least) forty-eight such transitions [6]. Perennial-to-annual transitions have been associated with the loss of awns in Danthonioideae and with increased above-ground biomass allocation in Pooideae [6,39]. The latter is hypothesised to have acted as an evolutionary precursor, facilitating multiple subsequent transitions to an annual life history within the clade possessing it (i.e. the ancestrally perennial core Pooideae; [6]). It is likely that this precursor pattern can be found in other clades of Poaceae as well, and that this contributes to the assumed life history lability in the family. In summary, previous studies have found numerous independent origins of annuals in grasses and, while this hints at evolutionary lability, we are unaware of any reports of annual-to-perennial reversals, or of any explicit tests of the assumed lability.

Thus, the main aim of this study is to compare four approaches for quantifying trait lability using life history evolution in Poaceae as an example. We discuss the merits and drawbacks of each approach and assess if they yield congruent results. The four approaches are: i) Estimates of phylogenetic signal using the D-statistic [40]; ii) Pattern of trait evolution inferred under an ancestral state reconstruction model that allows for transition rate heterogeneity (“hidden rates” model; [4]); Pattern of trait evolution inferred under an ancestral state reconstruction model, where transition rates depend on a simultaneous estimate of trait lability (threshold model; [26]); and iv) Comparison of observed transition rates to those expected by chance and under BM using simulations. The second aim of this study is to use model outcomes and inferences of lability to further understanding of life history variation and evolution in Poaceae. Thus, we frame our study around the following specific questions.

I. Can trait lability be determined using the four approaches tested and, if so, do they yield consistent results?
II. What is the phylogenetic distribution and mode of evolution of annuals in grasses?
III. Is life history a labile trait in grasses?

## Methods

### Life history and phylogenetic data

Life history data (annual/perennial) were assembled using GrassBase [41], filling in some gaps using the literature (Supplementary Table S1). Our full dataset included 11,539 species, of which 1989 are scored as annual (17.16%), 9434 as perennial (81.38%) and 116 as polymorphic. Exploratory analyses coding polymorphic species as either annual or perennial did not affect results, so polymorphic species were excluded from all final analyses, bringing the final total to 11,423 species (**Table 1**, Supplementary Data S1).

**Table 1.** Number and proportion of annual species in Poaceae overall and each subfamily separately for the full and phylogenetic datasets. Significant differences under the binomial test are shown in bold.

| Subfamily | Full dataset |  |  | Phylogenetic dataset |  |  | Binomial test |
| --- | --- | --- | --- | --- | --- | --- | --- |
|  | Species* | Annual species | Annuals % | Species | Annual species | Annuals % | P-value |
| Poaceae | 11 423 | 1 989 | 17.41 | 3 506 | 608 | 17.34 | 0.93 |
| Aristidoideae | 339 | 54 | 15.93 | 119 | 24 | 20.17 | 0.21 |
| Arundinoideae | 42 | 2 | 4.76 | 18 | 2 | 11.11 | 0.21 |
| Bambusoideae | 1 598 | 7 | 0.44 | 415 | 0 | 0.00 | 0.27 |
| Chloridoideae | 1 514 | 405 | 26.75 | 511 | 129 | 25.24 | 0.45 |
| Danthonioideae | 338 | 22 | 6.51 | 230 | 14 | 6.09 | 0.89 |
| Micrairoideae | 169 | 57 | 33.73 | 22 | 6 | 27.27 | 0.65 |
| Oryzoideae | 109 | 18 | 16.51 | 64 | 13 | 20.31 | 0.40 |
| Panicoideae | 3 317 | 961 | **28.97** | 820 | 204 | **24.88** | **<0.01** |
| Pooideae | 3 970 | 463 | **11.66** | 1 305 | 216 | **16.55** | **<0.0001** |

\*Species numbers from GrassBase [41].

To be able to perform comparative analyses and test for trait lability, phylogenetic information was obtained from Spriggs *et al*. [42]. This tree is based on fourteen nuclear and chloroplast subtrees grafted onto a chloroplast backbone, and contains 3595 species. We created a second dataset including only the species for which we had both life history and phylogenetic data, including 3506 species: 608 annuals (17.34%) and 2898 perennials (82.66%; Supplementary Data S2). To determine if the phylogenetic dataset is a good representation of the full dataset, we used a binomial test. We tested if the proportion of annuals differs significantly between the two datasets, for the Poaceae as a whole, as well as for each of six subfamilies analysed separately (see below).

### Testing trait lability using four approaches

Using the phylogenetic dataset, we tested for lability using four approaches: i) phylogenetic signal, ii) ancestral state reconstruction with the hidden rates model, iii) ancestral state reconstruction using the threshold model, and iv) simulations of expected transition rates under different scenarios. Two approaches (hidden rates and threshold models) allow illustration of the results over a phylogenetic tree, enabling visual determination of any differences among clades. The other two approaches (phylogenetic signal and simulations) provide only a single point estimate per analysis, limiting inference of any differences among clades. Given the long and probably complex evolutionary history of Poaceae [43], and to be able to separately model the presumed diverse histories of different major clades, we therefore performed these two analyses on six subfamilies separately, in addition to analysing the family as a whole: Aristidoideae, Chloridoideae, Danthonioideae, Oryzoideae, Panicoideae and Pooideae. These subfamilies were chosen because they include both annuals and perennials and our phylogenetic dataset includes enough species to robustly perform the analyses (>50 species; **Table 1**).

#### i) Phylogenetic signal

We calculated the D-statistic for Poaceae as a whole and each of the six subfamilies separately (see above) using ‘phylo.d’ in ‘caper’ in R (version 4.3.2; [40,44,45]). The D-statistic measures phylogenetic signal using the sum of sister-clade differences. D is expected to equal one when a trait is phylogenetically random, i.e., if trait variance is independent of phylogenetic relatedness. D is expected to equal zero under a BM model. Under this model, trait evolution is modelled as a “random walk”, where trait values shift over time and trait variance accumulates as lineages diverge [46]. For estimating the D statistic, the discrete trait is first modelled as continuous trait and then converted back to a discrete trait based on a threshold value to assign states (annual/perennial; [40]). The D estimated from the observed data is tested against simulations under D = 0 and D = 1. If D_observed_ = 0, the observed pattern fits expectations under BM, and we interpret this as stable. If D_observed_ = 1, the observed pattern fits the random scenario, and we interpret this as labile. If 0 < D_observed_ < 1 (i.e. significantly different from both 0 and 1), it is less labile and stable than expected, and we interpret this as intermediate.

#### ii) Ancestral state reconstruction with the hidden rates model

We reconstructed ancestral states using hidden Markov models (HMMs) that allow transition rates to vary over time and among clades, specifically, the “hidden rates model” of Beaulieu et al. [4]. The simplest model, the one-rate category model, has one rate of transitioning from perennial to annual and one rate from annual to perennial, i.e., a standard ancestral state reconstruction with all-rates-different (ARD; [47]). The model is then relaxed stepwise to add additional rate categories, such that each state (annual/perennial) may exist in one or more rate classes, representing different transition rates to and from observed states [4]. The two-rate category model thus allows a trait to transition in a “fast” and “slow” rate category, each with unique forward and reverse transition rates, as well as a rate of moving between rate categories. The three-rate category model includes “fast”, “medium” and “slow” rate categories, and this progresses in complexity up to the five-rate category model, the maximum currently implemented in ‘CorHMM’ [4,48]. The models do not allow for both the trait and transition rate category to change simultaneously but multiple shifts of any type can happen along a single branch. The rate of changing from one rate category to another is independent of the state (e.g., the rate of moving from rate category “fast” to “slow” is the same for state “perennial” and “annual”).

We tested the fit of models with one to five rate categories using maximum likelihood criteria and 100 random restarts, as implemented in ‘CorHMM’ [48], selecting the best fitting model using AICc values [49]. Then, we visualised reconstructed ancestral states based on the best-fitting model. Each life history state can exist in a “hidden state” based on its rate category; thus, possible ancestral states for the two-rate category model are perennial-fast, annual-fast, perennial-slow, annual-slow. We separately visualized ancestral states for life history (annual/perennial) and rate category (fast, slow, etc.) on different trees by summing the marginal probability of each state and rate at each node and labelling anything with a marginal probability < 0.75 as uncertain [48,50]. We interpreted clades with mainly “fast” rate categories as labile, and mainly “slow” as stable. Finally, to obtain an estimate of observed state changes, we summed those inferred between two directly connected (parent-descendent) nodes reconstructed with ≥0.75 certainty.

#### iii) Ancestral state reconstruction with the threshold model

The threshold model reconstructs the evolutionary history of a discrete trait using an underlying continuous variable, the liability factor, which determines the likelihood of a state change, in this case between perennial and annual [24,25]. We used the ‘ancThresh’ function in ‘phytools’ to estimate ancestral states and calculate liability factors, evolving under BM, through Bayesian Markov Chain Monte Carlo analysis [26,51], run for 5,000,000 generations with a 20% burnin and sampling every 1000 generations. We visualized ancestral states for life history (annual/perennial) on the tree labelling anything with a posterior probability (p.p.) < 0.95 as uncertain. Finally, to obtain an estimate of observed state changes, we summed those inferred between two directly connected (parent-descendent) nodes reconstructed with ≥0.95 certainty.

We visualised the liability factors in two ways. First, we calculated the mean liability factor at each node across all post-burnin runs and categorized them based on their absolute value: the third closest to the threshold were categorised labile; the third furthest from the threshold as stable; and the third with intermediate distance to the threshold as intermediate. Second, to further visualise how inferred life history lability varies among subfamilies, we created smoothed histograms of the liability factors across all post-burnin runs for each of the six subfamilies separately. We confirmed that the resulting patterns were not artefacts of summarising liability factors across runs by repeating both visualizations for ten randomly selected runs (they were not; results not shown).

#### iv) Simulations of transition rates

Lastly, we analysed perennial-to-annual (forward) and annual-to-perennial (reverse) transition rates separately. We compared both to those generated under a random scenario and under BM, for Poaceae as a whole and each of the six subfamilies separately. To do this, we estimated transition rates across the observed phylogenetic tree using ‘ace’ in ape [52]. We compared the fit of a model where forward and reverse shifts happen at the same rate (equal-rates, ER; [53]) to fit of a model where the two types of transitions may occur at different rates (ARD; [47]). For Poaceae as a whole and each the six subfamilies analysed separately, the ARD model was the best fit (not shown). This model was therefore used to estimate observed forward and reverse transition rates. We note that this model is unlikely to be the best overall (Supplementary Table 2), but since this test of lability required fitting 14,007 models (2001 traits across seven datasets), it was not feasible to test more complex models.

To compare observed transition rates to those expected under random and BM scenarios, we generated two datasets with 1000 simulated traits in each, for each of the seven clades. In the first, the random dataset, we randomly shuffled the observed data (annual/perennial) among the tips in the tree. In the second, the BM dataset, we simulated evolution of a continuous trait that varies between 0 and 1 under BM and using the observed tree and the function ‘BMplot’ in phytools [51]. The simulated data were converted to a binary trait by assigning the highest values as ‘annuals’, keeping the proportion of annuals as in the observed data for the Poaceae as a whole or each subfamily analysed separately. The remaining tips were assigned the state ‘perennial’. Transition rates across these simulated datasets were estimated using the ARD model as above.

Finally, we compared the transition rates inferred for observed data with those for simulated data. If a forward or reverse rate for observed data was not distinguishable from that for randomised data, we interpreted that transition to be labile. If the rate for observed data was not distinguishable from that simulated under BM, we interpreted that transition to be stable. If the rate for observed data was different to both random and BM scenarios, we interpreted that transition to be intermediate.

## Results

### The phylogenetic distribution of annual species in Poaceae

The distribution of annuals (17.41% of the species overall) differs greatly among subfamilies, from around a third of the species in some subfamilies to almost none in others (**table 1**). The highest percentages of annuals occur in the mainly (sub)tropical clades Micrairoideae (33.73%), Panicoideae (28.97%) and Chloridoideae (26.75%). Lower percentages occur in the largely temperate subfamilies Pooideae (11.66%) and Danthonioideae (6.51%), and in Arundinoideae (4.76%) and Bambusoideae (0.44%). The proportions of annuals in the full and phylogenetic datasets are mostly consistent; however, annuals are underrepresented in the phylogenetic dataset for Panicoideae and overrepresented for Pooideae (p<0.01; **Table 1**).

### Tests of trait lability

#### i) Phylogenetic signal

Life history variation in Aristidoideae and Pooideae has strong phylogenetic signal (not significantly different from D=0), while for Oryzoideae there is no phylogenetic signal (not significantly different from D=1). For Poaceae as a whole, Chloridoideae, Danthonioideae and Panicoideae the phylogenetic signal is intermediate (D=0.41, 0.45, 0.54 and 0.55, respectively), i.e., significantly different from both 0 and 1 (**table 2**).

**Table 2.** D-statistic estimated for Poaceae as a whole and six subfamilies separately. Significant differences are shown in bold. with the hidden rates model

| | D<br>(estimate) | p<br>($D=1$, random) | p<br>($D=0$, BM) |
| --- | --- | --- | --- |
| Poaceae | 0.406 | **0** | **0** |
| Aristidoideae | -0.069 | **0** | 0.613 |
| Chloridoideae | 0.420 | **0** | **0** |
| Danthonioideae | 0.538 | **0.003** | **0.047** |
| Oryzoideae | 0.904 | 0.285 | **0.002** |
| Panicoideae | 0.552 | **0** | **0** |
| Pooideae | 0.126 | **0** | 0.179 |

The model with three transition rate categories was the best fit (ΔAICc = 2.82 compared to the four-rate model, which had the second best fit; Supplementary Table S2). We identified a “slow”, “medium”, and “fast” rate category by summing forward and reverse transition rates within each rate category (**Table 3**). The distribution of these rate categories varies among clades (**Figure 1**). The ancestors of Bambusoideae and Pooideae are reconstructed with certainty as being in the “medium” rate category, but most other deep nodes are uncertain (e.g., the ancestors of Poaceae, BOP, PACMAD and the remaining subfamilies). In general, rates are higher in the PACMAD than the BOP clade: the “fast” rate category is found mainly in the PACMAD clade (specifically, Chloridoideae and Panicoideae), while deep nodes in Pooideae and Bambusoideae are in the “medium” category, with one large Pooideae clade transitioning to the “slow” category (core Pooideae, with *Loliinae* returning to the medium category).

**Table 3.**
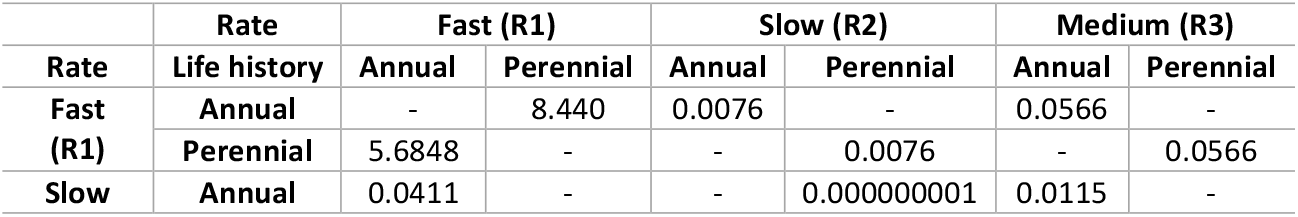
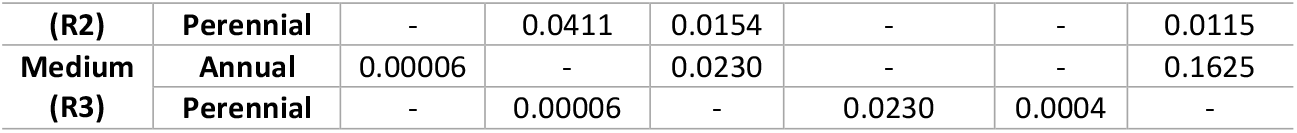
Rate matrix for best-fitting hidden rates model (three-rate model).

**Figure 1.**
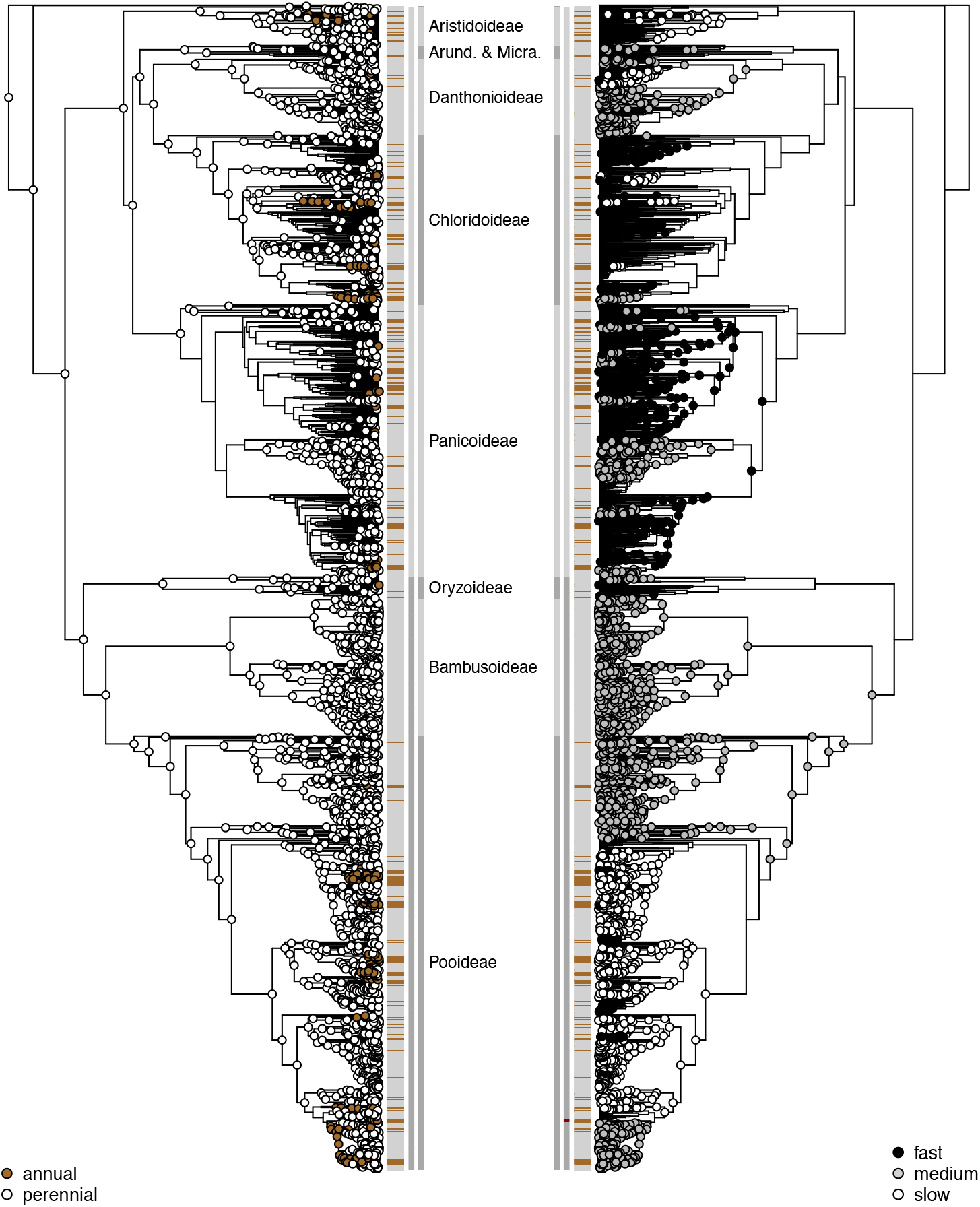
Ancestral state reconstruction for life history inferred from the best-fitting hidden rates model (Supplementary Table S2). Left: Ancestral states with a marginal probability ≥0.75 at each node, irrespective of rate category (white = perennial, brown = annual). Right: Ancestral rate categories with a marginal probability ≥0.75 at each node, irrespective of state (black = fast, grey = medium and white = slow). Nodes that are unlabelled (no filled circle) were equivocal (marginal probability <0.75 for state or rate). Tip states are shown as horizontal bars (grey = perennial, brown = annual). Outer grey vertical line: extent of labelled subfamilies (“Arund. & Micra.” = Arundinoideae and Micrairoideae). Inner grey vertical line: two major clades of grasses (dark grey = BOP [Bambusoideae, Oryzoideae, Pooideae]; light grey = PACMAD [Panicoideae, Aristidoideae, Chloridoideae, Micrairoideae, Arundinoideae, Danthonioideae]. The two successive sisters to the rest sampled here (*Puelia olyriformis* and *Pharus latifolius*) are not part of any labelled clade.

According to the best-fitting model, the ancestors of Poaceae, the major lineages BOP and PACMAD, and all subfamilies, are supported as being perennial (marginal likelihood ≥0.75; **Figure 1**). Annual species appear to have originated independently multiple times, even within individual subfamilies. Nodes inferred to have been ancestrally annual are relatively recent and found predominantly in Aristidoideae, Chloridoideae (e.g., *Muhlenbergia*), Panicoideae and Pooideae (e.g., *Bromus, Triticum, Loliinae*). Overall, we identified at least 63 transitions from perennial to annual and at least 13 reversals from annual to perennial, based only on nodes with ≥0.75 support.

#### iii) Ancestral state reconstruction with the threshold model

The liability factor estimated at each node ranged from -16.48 to 29.78 across the post-burnin models (Supplementary Figure S1), with a threshold of zero signifying a state transition (positive values indicate perennial; negative values indicate annual). The per-node absolute means calculated across the posterior sample ranged from 0.001 to 13.18, with high values (far from the threshold, indicating stability) found at the root and all of the deepest nodes, including the ancestors of all subfamilies (**Figure 2**). Lower values (close to the threshold, indicating lability) were found towards the tips of the tree, across all the major clades, apart from Bambusoideae, which stands out as predominantly stable. The smoothed histograms reveal a high degree of similarity in estimated liability factors among the six subfamilies analysed separately, displaying only slight variation in the position and height of the peaks (Supplementary Figure S2). Notably, while most subfamilies exhibit a single prominent peak in the perennial state, not far from the threshold, the Oryzoideae stands out with multiple peaks close to the threshold or in the perennial state, indicating a more complex pattern of evolutionary state change within this group.

**Figure 2.**
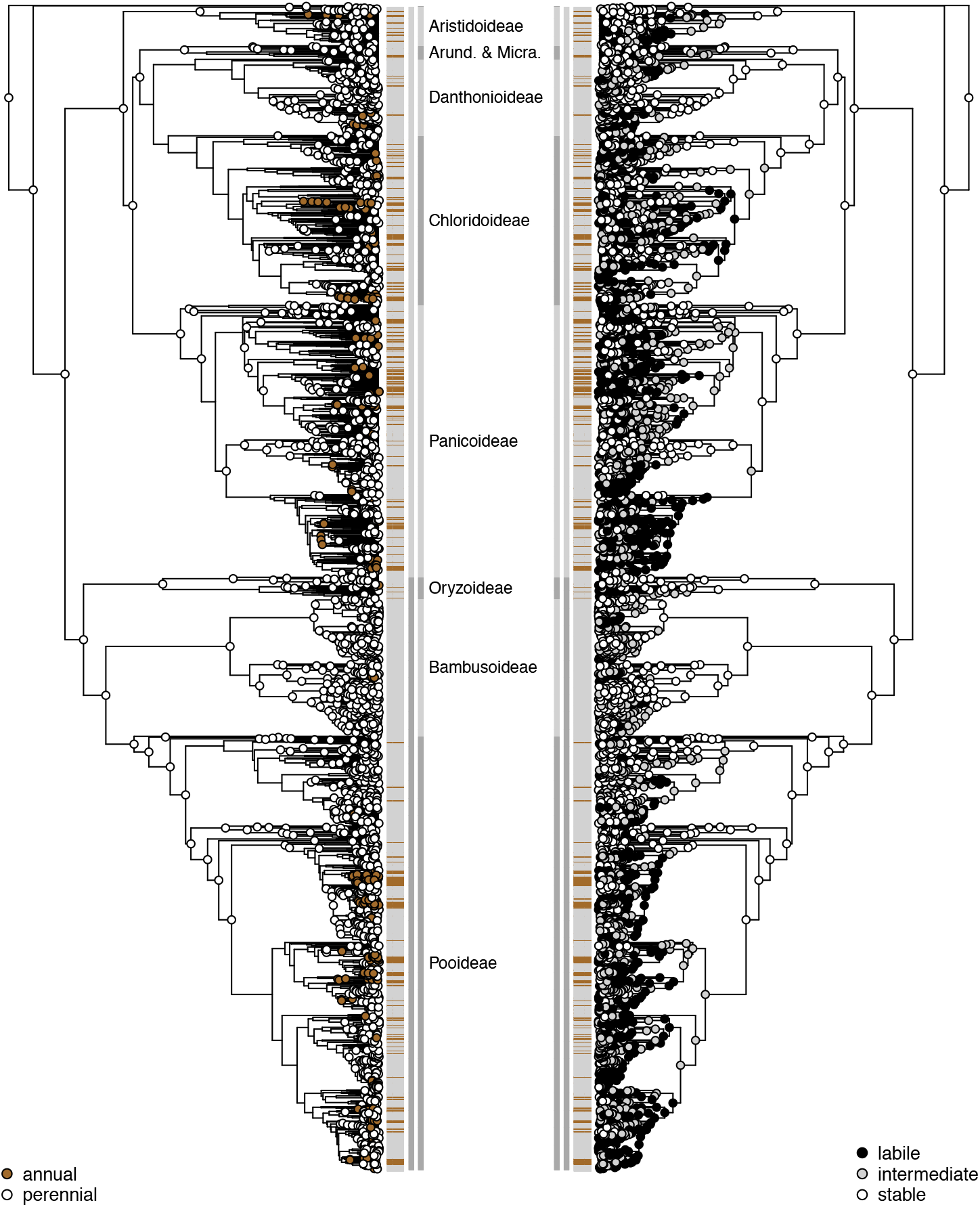
Ancestral state reconstruction for life history inferred from the threshold model. Left: Ancestral states reconstructed with p.p.≥0.95 at each node, irrespective of liability factor (white = perennial, brown = annual). Unlabelled nodes (no filled circle) were equivocal (p.p.<0.95). Right: Mean absolute liability factor at each node, categorized as labile (closest to the threshold), intermediate, and stable (furthest from the threshold). Tip states are shown as horizontal bars (grey = perennial, brown = annual). Outer grey vertical line: extent of labelled subfamilies (“Arund. & Micra.” = Arundinoideae and Micrairoideae). Inner grey vertical line: the two major clades of grasses (dark grey = BOP [Bambusoideae, Oryzoideae, Pooideae]; light grey = PACMAD [Panicoideae, Aristidoideae, Chloridoideae, Micrairoideae, Arundinoideae, Danthonioideae]. The two successive sisters to the rest sampled here (*Puelia olyriformis* and *Pharus latifolius*) are not part of any labelled clade.

The ancestral state reconstruction indicates that the ancestors of Poaceae, the major lineages BOP and PACMAD, and all subfamilies except Danthonioideae are perennial (p.p.≥0.95; **Figure 2**). Annuals appear to have originated independently multiple times, even within individual subfamilies. Nodes inferred to have been ancestrally annual are relatively recent and found mainly in Aristidoideae, Danthonioideae, Chloridoideae (e.g., *Muhlenbergia*), Panicoideae and Pooideae (e.g., *Bromus, Triticum, Loliinae*). We identified at least 19 transitions from perennial to annual and at least 32 reversals from annual to perennial, based only on nodes with p.p.≥0.95).

#### iv) Simulations of transition rates

As expected, transition rates estimated for the randomised data are higher than those estimated from the data simulated under BM (**Figure 3**). Observed transition rates for Poaceae are intermediate to those for randomised data and those simulated under BM, transition rates for Pooideae are similar to those simulated under BM, and transition rates for Oryzoideae, Panicoideae, Chloridoideae and Danthonioideae overlap with or are higher than for randomised data. Finally, we note that forward transitions occur at lower rates than reverse across all datasets, most likely an artefact of the simple model used.

**Figure 3.**
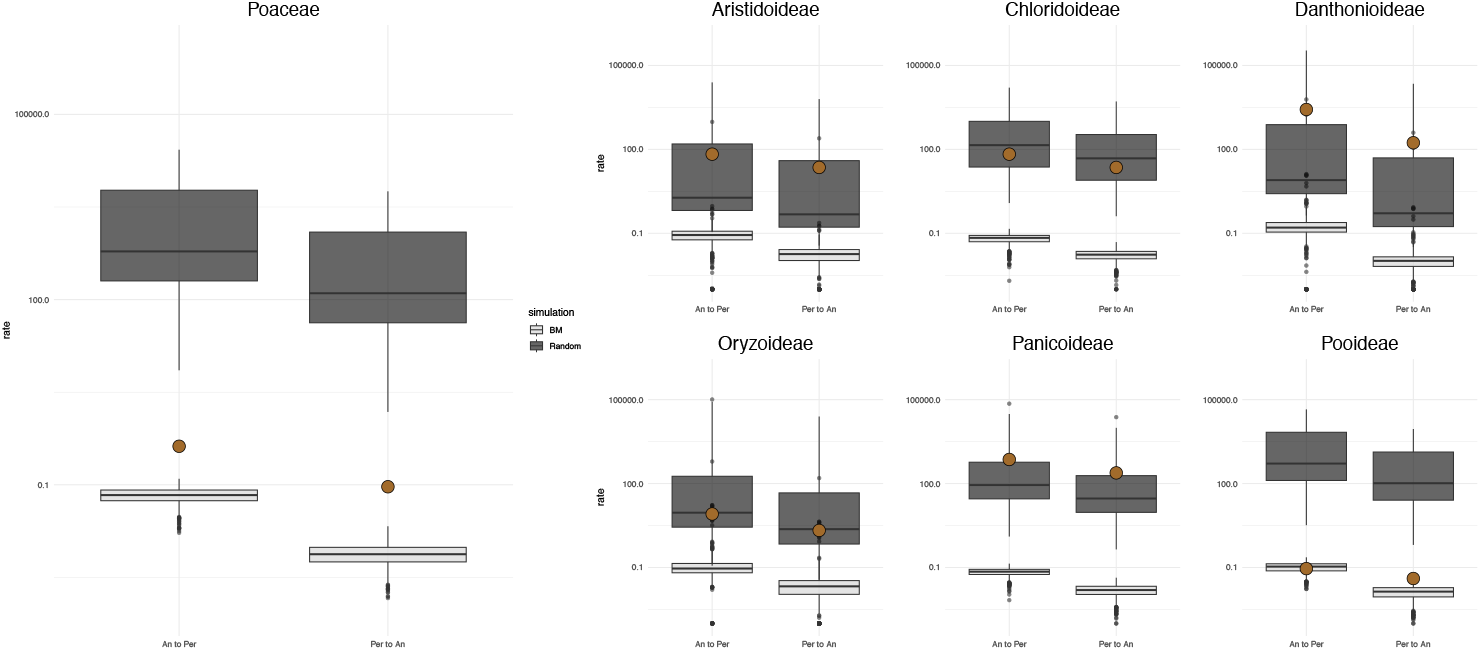
Distribution of transition rates for observed (filled circles), randomised data (dark grey) and data simulated under Brownian motion (BM, light grey) estimated the all-rates-different (ARD) model. Outliers are displayed as dark grey (random) or light grey (BM) dots outside the boxplots.

## Discussion

### Comparison of trait lability estimates using four different approaches

Trait lability has been tested using a range of approaches in the literature [11,12,21]. We tested four commonly used ones to analyse life history variation in grasses (Poaceae) and found that they differed greatly in their inferences of trait lability: no two approaches consistently gave the same results (**table 4**). The results of the hidden rates model [4] and simulations were the most similar, agreeing on four inferences, perhaps because they are both based on transition rates. The second most similar results were for the D-statistic [40] and hidden rates model, and D-statistic and simulations, with three shared results each. The threshold model [26] was the least consistent with the other approaches, yielding the same result as another only in one case (the hidden rates model for Poaceae as a whole, and the result was inconclusive). In all, there were 11 shared across 28 tests performed.

**Table 4.** Summary of the four approaches used to test for trait lability and main results for each.

| Table 4. Summary of the four approaches used to test for trait liability and main results for each. |  |  |  |  |
| --- | --- | --- | --- | --- |
| Approach | Phylogenetic signal | Ancestral state reconstruction (hidden rates model) | Ancestral state reconstruction (threshold model) | Simulations of transition rates |
| Type of test | Objective | Interpretive | Interpretive | Objective |
| Description of test | Compares D-statistic to random (labile) and Brownian Motion (stable). | Determines transition rate category at each node from fast (labile) to slow (stable). | Calculates liability factor, determining chance of a state change, from liable (labile) to less liable (stable). | Compares observed transition rates to random (labile) and Brownian Motion (stable). |
| Per-clade results*: |  |  |  |  |
| Poaceae | intermediate | variable | variable | intermediate |
| Aristidoideae | stable | stable | variable | labile |
| Chloridoideae | intermediate | labile | variable | labile |
| Danthonioideae | intermediate | stable | variable | labile |
| Oryzoideae | labile | labile | variable | labile |
| Panicoideae | intermediate | labile | variable | labile |
| Pooideae | stable | stable | variable | stable |

\*Key: stable= trait clearly stable, labile= trait clearly labile, variable= trait both stable and labile, intermediate= trait intermediate to labile and stable.

The level of inconsistency also differed among the clades analysed. The most conflicting results were for the Danthonioideae, with each approach suggesting a different inference of trait lability (intermediate, stable, variable, labile; **Table 4**). The most consistent results were for Poaceae (intermediate or variable), Oryzoideae (labile or variable) and Pooideae (stable or variable). For Aristidoideae, results of the D-statistic and the hidden rates model both suggested trait stability – but trait lability was inferred based on the simulations. Thus, even if two of the approaches agreed, both lability and stability were inferred, making results for this clade highly inconsistent.

The four approaches differ their objectiveness (objective/interpretive). Phylogenetic signal and the simulations are the two most objective approaches, as they compare observed results to patterns expected for trait lability or stability. Using these two approaches we were able to infer life history as labile, stable or intermediate for all clades (**Table 4**). However, these approaches provide only a single estimate per clade, so may obscure any within-clade nuances. The ancestral state reconstructions allow different interpretations for different parts of a clade, but have the drawback that visual inspection of the estimated transition rates (hidden rates) or liability (threshold) is necessary, which can reduce the objectivity of these tests. Nevertheless, we were able to infer evolutionary lability or stability for all six subfamilies analysed separately using the hidden rates model, with only Poaceae as a whole being inferred as ‘variable’, which is perhaps unsurprising given the size of the clade. In contrast, we did not obtain a clear result for any of the clades using the threshold analysis (**Table 4**), despite this analysis being formulated around a parameter determining a trait’s liability to change [26]. Instead, the liability factor itself appears to be a highly labile trait, switching between labile and stable values numerous times throughout the tree (**Figure 2**). Nevertheless, this approach has been used to clearly define trait lability before [11], and the inconclusiveness of our results may be due to the size of the clade analysed (Poaceae as a whole), coupled with high transition rates in certain parts of it (**Figures 2, 3**).

The objective and interpretative approaches revealed different aspects of the same data and may be appropriate for different purposes. The objective approaches are easy to interpret and may be appropriate for large-scale analyses of many clades or traits, where it is not possible to dive into within-clade differences. However, the interpretative approaches have revealed that lability itself is not constant, but acts as an independent trait with its own evolutionary history. Such within-clade nuances may reveal additional traits or aspects of the evolutionary history that could influence lability (e.g. precursor traits; [3,4,6]), thus deepening both the evidence for and understanding of the lability of the trait of interest. Ultimately, which approach is most appropriate will depend on the exact question, but if the aim is to determine trait lability, comparing observed state transition rates to expectations under models of trait stability and lability will provide a test that is both objective and based on explicit criteria. Despite longstanding interest in identifying labile traits or comparing trait lability among clades (e.g. [10–12,15,17]), this approach seems to have been little used.

Importantly, our results highlight that choice of method really matters, raising the question whether our results are particular to life history evolution in grasses or more general, being true for other traits in grasses (e.g., salt tolerance or C_4_ photosynthesis [5,21]) or other groups (e.g. *Solanum* or Montiaceae [11,12]), where trait lability has previously been tested using one or more of the approaches compared here. Certainly, our findings question the comparability of previous estimates of trait lability, specifically, whether they allow drawing general conclusions about the evolutionary flexibility of certain traits, or whether conclusions must be conditioned on the trait, plant group and approach in question.

### Distribution and evolution of annuals in grasses

We find stark variation in how life history is distributed and evolves across grasses. Annual species occur in all subfamilies but are unevenly distributed among them (**Figure 1, Table 1**). Most annuals are found in the mainly (sub)tropical PACMAD clade (75%), with as much as a third of the species being annual in some subfamilies (Micrairoideae, 34%; Panicoideae, 29% and Chloridoideae 27%). These clades include both C_3_ and C_4_ species [54]. In contrast, lower proportions of annuals are found in the temperate, cooler-climate BOP clades, comprising entirely C_3_ species (Pooideae, 12% and Bambusoideae, 0.44%), and even the C_3_ temperate PACMAD clade Danthonioideae (7%). All bamboos, save the seven annual species of *Raddiella*, are perennial, which conforms to their status as the only woody clade in the grasses [55]. The uneven distribution of annuals among clades may reflect their large variation in morphology, growth forms and habitats, a result of over 100 Ma of evolution [56–58]. The finding that annual grasses are concentrated in PACMAD is new and suggests that C_4_ photosynthesis and an annual life history strategy are favoured in the same types of environments (hot and dry; e.g. [30,33,38,59]). Further research is needed to determine whether PACMAD annuals more likely utilise the C_3_ or C_4_ photosynthetic pathway and whether there is any ecological or evolutionary correlation between these traits.

Grasses were most likely ancestrally perennial [6,38,39] and, accordingly, the ancestors of Poaceae, the major clades BOP and PACMAD, and all subfamilies were reconstructed as perennial with certainty under both the hidden rates and threshold models (**Figures 1, 2**). No deep nodes were reconstructed as annual with certainty, only more recent ones. We estimated 63 origins of annuals under the hidden rates model and 19 under the threshold model (Supplementary Text S1). These numbers are certainly underestimates, partly due to uncertainty in the reconstruction in some parts of the tree and limited taxon sampling. For example, a previous study estimated 48 origins in Pooideae alone [6]. However, not all reconstructed origins are likely to be “true”. Based on the threshold model, there is at least one origin of annuality that is directly followed by reversals to perenniality in Bambusoideae. It is highly doubtful that this represents the true evolutionary history in the bamboos, and more likely it is caused by a “wobble” of the liability factor. The same may be true for other reconstructed state transitions. Thus, reconstructions of this nature can both under- and overestimate true evolutionary events.

Nevertheless, it is still interesting to note that both the hidden rates and threshold models inferred a number of annual-to-perennial reversals (13 and 32, respectively). Such reversals have traditionally been controversial, not least because they are thought to be very rare in nature, although the number of examples is accumulating across flowering plants [11,38,60,61]. We consider potential “reversals” to be new gains of perennialism, in accordance with Dollo’s law [62], and given widespread interest in perennialising grain crops [63,64], it is relevant to pinpoint where such secondary gains of perennialism may naturally have occurred. Determining this in detail would require more fine-grained analyses, but our reconstructions provide a starting point: both ancestral state reconstructions find reversals in the *Loliinae* (Pooideae) and in *Muhlenbergia* subg. *Muhlenbergia* (*sensu* Peterson [65]; Chloridoideae). These clades are mostly frost tolerant but generally occur in areas without severe winters [50], which agrees with the reputation of annuals prevailing in warm temperate climates [11,33,66]. The *Loliinae* consists of annuals, long-lived perennials and short-lived perennials that may persist and flower in multiple seasons under favourable conditions (*Lolium westerwoldicum*; [67]). Life history shifts may be more likely in such clades with “intermediate” forms, such as or short-lived, non-persistent perennials, or with resource allocation patterns similar to annuals [4,6,30]. The same is unlikely to be happening in *Muhlenbergia* subg. *Muhlenbergia*, however, as these are rhizomatous, C4 perennials [68].

### Is life history a labile trait in grasses?

Life history has long been considered one of the most labile traits in plants [19,29], probably being more labile than in animals [32]. Grasses comprise more annual species than almost any other group of plants [30], but does this make life history a labile trait in grasses? Overall, our results show that the answer is: not necessarily. Instead, trait lability itself varies among clades and over time and inferences of lability are highly contingent on the approach used to test for it. If we disregard the results of the threshold analyses (which were inconclusive, and see Supplementary Text S1), we find that life history evolution is definitely labile in only three clades (Chloridoideae, Panicoideae and Oryzoideae) and stable in one clade (Pooideae; **Table 4**). The results for Aristidoideae and Danthonioideae were less conclusive. According to the simulations, state transitions in both clades conformed to expectations under the random scenario (**Figure 3**), i.e. trait lability. According to the hidden rates model, however, transition rates are lower in these two clades than other clades, suggesting trait stability (**Figure 1, Table 4**). Given that the latter result only holds relative to other clades, we favour the results of the simulations, which hold relative to expectations specific to each clade. These suggest that life history evolution is labile in all clades of grasses, except Pooideae (Supplementary Text S1). We were (consistently) not able to determine life history lability across grasses as a whole and the simulations suggest intermediate lability (**Figure 3**), which may be unsurprising for a large, old clade like the grasses. This cautions against trying to find generalities in the evolution of complex trait syndrome across diverse, old and globally distributed clades (cf. [4,5]).

## Conclusions

Our study highlights the complexity of assessing trait lability. We found that different approaches give varying conclusions, no two methods consistently gave the same result and no single approach allowed unambiguously determining life history as labile or stable across all clades. Overall, lability itself is labile, and a relative concept that only gains meaning through comparative analysis. The chosen approach should depend on the exact question, but only comparison to expectations under models of trait stability and lability will provide an objective test based on explicit criteria. Our study further demonstrates that most annual grasses are concentrated in clades that occupy warm climates and include both C_3_ and C_4_ species. The nature of any potential ecological and evolutionary link between the annual life history strategy and C_4_ photosynthesis warrants further research. Annuals are inferred to have originated frequently and recently–but we also identify several potential reversals to perenniality. These may provide starting points for more fine-grained analyses in support of initiatives to perennialise food crops. Finally, and contrary to expectations based on the numerous and widespread life history transitions across the grasses, we show that life history is not a labile trait throughout the grasses, but stable in the largest clade, the Pooideae, comprising crops such as wheat and barley. Our findings caution against broad generalizations across diverse, old, and widely distributed clades and highlight the importance of objective tests of trait lability.

## Author contributions

L.S. and A.M.H. designed the study. L.S. compiled and analysed the data. L.S. and A.M.H. interpreted the results and wrote the paper.

## Acknowledgements

We sincerely thank Marc Fradera-Soler and Siri Fjellheim for insightful discussions and Maria (Bat) Vorontsova for providing (via GrassBase) and checking the life history data.

