## Supplementary Material for "Testing for trait lability at macroevolutionary scales: is life history evolution labile in grasses?"

#### Contents

**Table S1**. Synonyms and life history data for species not included in GrassBase.

**Table S2**. Log-likelihoods and AICc values of HMM models with different numbers of rate categories.

**Figure S1**. Convergence plot for the ancestral state reconstruction using the threshold model.

**Figure S2**. Liability factors estimated using the threshold model.

**Text S1**. Supplementary Discussion

**Data S1**. The full life history dataset analysed. (Provided as a separate .csv file.)

**Data S2**. The reduced (phylogenetic) life history dataset analysed. (Provided as a separate .csv file.)

| **Table S1.** Synonyms and life history data for species in the phylogenetic dataset but not in GrassBase [1] Synonyms were checked in POWO* (https://powo.science.kew.org/) and WFO° (http://www.worldfloraonline.org/), and alternative names matched to GrassBase for life history data, unless otherwise specified (* = POWO, ° = WFO). | | |
|---|---|---|
| **Species in tree** | **Synonym of:** | **Life History:** |
| *Alloteropsis semialata* subsp. *eckloniana* | Accepted name* | Perennial° |
| *Alloteropsis semialata* subsp. *semialata* | Accepted name* | Perennial° |
| *Anthenantia rufa* | Accepted name* | Perennial* |
| *Apocopis collinus* | Accepted name* | Perennial* |
| *Apocopis intermedius* | Accepted name* | Perennial* |
| *Arundinaria tecta* | Accepted name* | Perennial |
| *Aulonemia clarkiae* | *Olmeca clarkiae** | Perennial |
| *Aulonemia fulgor* | *Olmeca fulgor** | Perennial |
| *Austrofestuca triticoides* | *Poa triticoides** | Perennial*, collected under *Austrofestuca littoralis* |
| *Avena insularis* | Accepted name*° | Annual* |
| *Bambusa moreheadiana* | *Mullerochloa moreheadiana** | Perennial |
| *Brachypodium arbuscula* | Accepted name* | Perennial* |
| *Brachypodium retusum* | Accepted name* | Perennial*, collected under *Brachypodium plukenetii* |
| *Briza poimorpha* | *Chascolytrum poimorphum** | Perennial*, collected under *Microbriza poimorpha* |
| *Bromus korotkiji* | *Bromus pumpellianus** | Perennial* |
| *Cenchrus chilensis* | Accepted name* | Perennial*, collected under *Pennisetum chilense* |
| *Cenchrus incertus* | *Cenchrus spinifex** | Annual |
| *Centrochloa singularis* | *Axonopus singularis** | Annual* |
| *Chasmanthium sessiliflorum* | *Chasmanthium laxum** | Perennial |
| *Chusquea aff fendleri* | Possibly Ch*usquea fendleri.* Genus is perennial*° | Perennial*° |
| *Coelachne simpliciuscula* | Accepted name* | Annual° |
| *Cortaderia archboldii* | *Chimaerochloa archboldii** | Perennial |
| *Cortaderia fulvida* | *Austroderia fulvida** | Perennial |
| *Cortaderia richardii* | *Austroderia richardii** | Perennial |
| *Cortaderia splendens* | *Austroderia splendens** | Perennial |
| *Cortaderia toetoe* | *Austroderia toetoe** | Perennial |
| *Cortaderia turbaria* | *Austroderia turbaria** | Perennial |
| *Distichlis eludens* | Accepted name* | Perennial*, collected under *Reederochloa eludens* |
| *Drepanostachyum sengteeanum* | Genus is perennial* | Perennial* |
| *Echinochloa crus-galli* | Accepted name* | Annual* |
| *Elymus burchan-buddae* | Accepted name* | Perennial* |
| *Elymus enysii* | *Stenostachys enysii** | Perennial |
| *Elymus multiflorus* | *Anthosachne kingiana* subsp. *multiflora** | Perennial*, collected under *Anthosachne kingiana* |
| *Elymus reflexiaristatus* | *Pseudoroegneria reflexiaristata** | Perennial*, collected under Elymus aegilopoides |
| *Elymus scaber* | Anthosachne scabra* | Perennial*, collected under Elymus *scaber* |
| *Elymus tangutorum* | *Elymus dahuricus** | Perennial |
| *Elymus virginicus* var. *virginicus* | *Elymus virginicus°* | Perennial |
| *Eragrostis obtusiflora* | *Kalinia obtusiflora** | Perennial |
| *Fargesia frigidis* | Accepted name* | Perennial* |
| *Festuca borderii* | *Festuca borderei* | Perennial |

| | | |
|---|---|---|
| *Festuca coerulescens* | *Patzkea coerulescens** | Perennial*, collected under *Festuca coerulescens* |
| *Festuca matthewsii* subsp. *aquilonia* | *Festuca matthewsii** | Perennial* |
| *Festuca pseudovina* | *Festuca pulchra**° | Perennial*° |
| *Festuca rivas martinezii* subsp. *rectifolia* | *Festuca lambinonii** | Perennial |
| *Gigantochloa verticillata* | Accepted name* | Perennial* |
| *Greslania circinata* | Accepted name* | Perennial* |
| *Helictochloa armeniaca* | Accepted name* | Perennial*, collected under *Helictotrichon armeniacum* |
| *Helictochloa blaui* | Accepted name* | Perennial*, collected under *Helictotrichon blaui* |
| *Helictochloa pratensis* | Accepted name* | Perennial*, collected under *Helictotrichon pratense* |
| *Helictotrichon krischae* | Genus is perennial*° *Helictotrichon x krischae** | Perennial*° |
| *Helictotrichon marginatum* | *Helictochloa marginata** | Perennial*, collected under *Helictotrichon occidentale* |
| *Helictotrichon setaceum* subsp. *setaceum* | *Helictotrichon setaceum** | Perennial* |
| *Heterachne abortivum* | *Heterachne abortiva* | Annual |
| *Himalayacalamus intermedius* | Genus is perennial*° | Perennial*° |
| *Hystrix californica* | *Elymus californicus** | Perennial*, collected under *Hystrix californica* |
| *Hystrix coreana* | *Elymus coreanus** | Perennial*, collected under *Hystrix coreana* |
| *Hystrix duthiei* | *Leymus duthiei** | Perennial |
| *Hystrix gracilis* | *Stenostachys gracilis** | Perennial*, collected under *Hystrix gracilis* |
| *Hystrix komarovii* | *Leymus komarovii** | Perennial*, collected under *Hystrix komarovii* |
| *Hystrix laevis* | *Stenostachys laevis** | Perennial*, collected under *Hystrix laevis* |
| *Hystrix patula* | *Elymus hystrix** | Perennial |
| *Indocalamus hamadae* | *Indocalamus tessellatus** | Perennial |
| *Kinabaluchloa ridleyi* | Genus is perennial*° | Perennial*° |
| *Koeleria aff novozelandica* | Assuming *Koeleria novozelandica** | Perennial |
| *Koeleria digorica* | Genus is perennial*° | Perennial*° |
| *Lasiacis sorghoidea* | *Lasiacis maculate** | Perennial |
| *Leymus ajanensis* | Accepted name* | Perennial* |
| *Melica sabinei* | *Pleuropogon sabinei** | Perennial |
| *Miscanthus giganteus* | Accepted name* | Perennial* |
| *Muhlenbergia brandegeei* | *Muhlenbergia brandegeei* | Annual |
| *Muhlenbergia bryophilus* | Accepted name* | Annual*, collected under *Aegopogon bryophilus* |
| *Muhlenbergia cenchroides* | Accepted name* | Annual*, collected under *Aegopogon cenchroides* |
| *Muhlenbergia geminiflora* | *Muhlenbergia cenchroides** | Annual |
| *Muhlenbergia multiflora* | Accepted name* | Annual*, collected under *Redfieldia flexuosa* |
| *Muhlenbergia plumiseta* | Accepted name* | Annual*, collected under *Pereilema ciliatum* |
| *Muhlenbergia spatha* | Accepted name* | Annual*, collected under *Schaffnerella gracilis* |

| | | |
|---|---|---|
| *Muhlenbergia tricholepis* | Accepted name* | Perennial*, collected under *Blepharoneuron tricholepis* |
| *Neomicrocalamus yunnanensis* | *Melocalamus yunnanensis** | Perennial |
| *Neurolepis aperta* | *Chusquea spectabilis** | Perennial |
| *Neurolepis aristata* | *Chusquea aristate** | Perennial |
| *Neurolepis asymmetrica* | *Chusquea asymmetrica** | Perennial |
| *Neurolepis elata* | *Chusquea elata** | Perennial |
| *Neurolepis nana* | *Chusquea nana** | Perennial |
| *Neurolepis pittieri* | *Chusquea magnifolia** | Perennial |
| *Neurolepis rigida* | *Chusquea rigida** | Perennial |
| *Neurolepis villosa* | *Chusquea villosa** | Perennial |
| *Neurolepis virgata* | *Chusquea cylindrica** | Perennial |
| *Oncorachis ramosa* | Accepted name* | Perennial*, collected under *Streptostachys ramosa* |
| *Ophiochloa hydrolithica* | *Axonopus hydrolithicus** | Perennial |
| *Oryza glumipatula* | *Oryza rufipogon*° | Perennial |
| *Oryzopsis aequiglumis* | *Piptatherum aequiglume*° | Perennial |
| *Oryzopsis alpestris* | *Piptatherum alpestre** | Perennial |
| *Oryzopsis angustifolia* | *Piptatherum angustifolium** | Perennial |
| *Oryzopsis canadensis* | *Piptatherum canadensis** | Perennial |
| *Oryzopsis chinensis* | Accepted name* | Perennial |
| *Oryzopsis coerulescens* | *Piptatherum coerulescens** | Perennial |
| *Oryzopsis ferganensis* | *Piptatherum ferganensis** | Perennial |
| *Oryzopsis gracilis* | *Piptatherum gracile** | Perennial |
| *Oryzopsis hilariae* | *Piptatherum hilariae** | Perennial |
| *Oryzopsis holciformis* | *Piptatherum holciforme** | Perennial |
| *Oryzopsis lateralis* | *Piptatherum laterale** | Perennial |
| *Oryzopsis latifolia* | *Piptatherum latifolium** | Perennial |
| *Oryzopsis micrantha* | *Piptatherum micranthum*° | Perennial |
| *Oryzopsis miliacea* | *Piptatherum miliaceum** | Perennial |
| *Oryzopsis molinioides* | *Piptatherum molinioides** | Perennial |
| *Oryzopsis munroi* | *Piptatherum munroi** | Perennial |
| *Oryzopsis obtusa* | *Piptatherum obtuse** | Perennial*, collected under *Piptatherum kuoi* |
| *Oryzopsis paradoxa* | *Achnatherum paradoxum** | Perennial*, collected under *Piptatherum paradoxum* |
| *Oryzopsis sogdiana* | *Piptatherum sogdianum** | Perennial |
| *Oryzopsis songarica* | *Piptatherum songaricum** | Perennial |
| *Oryzopsis sphacelata* | *Piptatherum sphacelatum**<br>Genus *Piptatherum* is perennial*° | Perennial*° |
| *Oryzopsis vicaria* | *Piptatherum microcarpum** | Perennial |
| *Oryzopsis virescens* | *Achnatherum virescens** | Perennial*, collected under *Piptatherum virescens* |
| *Oryzopsis webberi* | *Eriocoma webberi** | Perennial*, collected under *Stipa webberi* |
| *Panicum discolor* | *Panicum penicillate** | Perennial*, collected under *Panicum penicillatum* |
| *Pappostipa chrysophylla* | Accepted name* | Perennial*, collected under *Stipa chrysophylla* |
| *Pappostipa vaginata* | Accepted name* | Perennial*, collected under *Stipa vaginata* |
| *Paspalum juergensii* | Accepted name* | Perennial* |
| *Paspalum trichotomum* | Accepted name* | Perennial* |
| *Pentameris dregeana* | Accepted name* | Perennial*, collected under *Pentameris distichophylla* |
| *Pentameris galipinii* | *Pentameris galpinii** | Perennial |

| | | |
|---|---|---|
| *Pentameris heptameris* | *Pentameris heptamera°* | Perennial |
| *Pentameris juncifolia* | *Pentameris eriostoma** | Perennial |
| *Pentameris minor* | Accepted name* | Perennial* |
| *Pleioblastus fortunei* | *Pleioblastus variegatus** | Perennial*, collected under *Pleioblastus fortunei* |
| *Pleioblastus hsienchuensis* | Accepted name* | Perennial*, collected under *Sinobambusa seminuda* |
| *Poa cockayneana* | Accepted name* | Perennial* |
| *Poa hartzii* subsp. *hartzii* | *Poa hartzii°* | Perennial |
| *Poa violacea* subsp. *aetnensis* | If *Poa violacea* then *Bellardiochloa variegate** | Perennial*, collected under *Poa variegata* |
| *Pogoneura biflora* | *Pogononeura biflora* | Annual |
| *Psathyrostachys rupestris* subsp. *rupestris* | *Psathyrostachys rupestris°* | Perennial |
| *Rostraria villosa* | Genus is annual*° | Annual*° |
| *Rupichloa acuminata* | Accepted name* | Perennial*, collected under *Streptostachys acuminata* |
| *Rytidosperma aureocephalum* | possibly *Tenaxia aureocephalum** | Perennial*° |
| *Rytidosperma cf corinum* | *Rytidosperma corinum** | Perennial |
| *Rytidosperma curvum* | Genus is perennial*° | Perennial*° |
| *Rytidosperma davyi* | *Merxmuellera davyi** | Perennial |
| *Rytidosperma distichum* | *Tenaxia disticha** | Perennial |
| *Rytidosperma drakensbergense* | Genus is perennial*° | Perennial |
| *Rytidosperma durum* | possibly *Tenaxia dura** Both genera are perennial*° | Perennial*° |
| *Rytidosperma guillarmodiae* | possibly *Tenaxia guillarmodiae?* Both genera are perennial*° | Perennial |
| *Rytidosperma macowanii* | Genus is perennial*° | Perennial |
| *Rytidosperma pictum* var. *pictum* | *Rytidosperma pictum°* | Perennial |
| *Rytidosperma purpureum* | Genus is perennial*° | Perennial |
| *Rytidosperma rufum* | Genus is perennial*° | Perennial |
| *Rytidosperma schismoides* | Genus is perennial*° | Perennial |
| *Rytidosperma stereophyllum* | Genus is perennial*° | Perennial |
| *Rytidosperma strictum* | possibly *Tenaxia stricta* | Perennial*° |
| *Rytidosperma subulatum* | *Tenaxia subulate** | Perennial |
| *Rytidosperma telmaticum* | Genus is perennial*° | Perennial*° |
| *Rytidosperma tenellum* | Genus is perennial*° | Perennial*° |
| *Saxipoa saxicola* | Accepted name* | Perennial* |
| *Schismus pleuropogon* | *Tribolium pleuropogon** | Perennial |
| *Secale strictum* subsp. *strictum* | *Secale strictum** | Perennial*, collected under *Secale montanum* |
| *Semiarundinaria makinoi* | Genus is perennial*° | Perennial |
| *Shibataea kumasaca* | *Shibataea kumasasa* | Perennial |
| *Stipa arcaensis* | *Nassella arcaensis** | Perennial |
| *Stipa arcuata* | *Nassella arcuate** | Perennial |
| *Stipa argentinensis* | *Nassella argentinensis** | Perennial |
| *Stipa brachychaetoides* | *Nassella brachychaetoides** | Perennial |
| *Stipa brachyphylla* | *Nassella brachyphylla** | Perennial |
| *Stipa charruana* | *Nassella charruana** | Perennial |
| *Stipa clarazii* | *Nassella longiglumis** | Perennial |
| *Stipa cordobensis* | *Nassella cordobensis** | Perennial |
| *Stipa curamalalensis* | *Nassella curamalalensis** | Perennial |
| *Stipa curtiseta* | *Stipa spartea°* | Perennial |
| *Stipa depauperata* | *Nassella depauperate** | Perennial |
| *Stipa filiculmis* | *Nassella filiculmis** | Perennial |

| | | |
|---|---|---|
| *Stipa hyalina* | *Nassella hyaline** | Perennial |
| *Stipa inconspicua* | *Nassella inconspicua** | Perennial |
| *Stipa leucotricha* | *Nassella leucotricha** | Perennial |
| *Stipa lobata* | *Eriocoma lobata** or *Stipa robusta*° | Perennial |
| *Stipa manicata* | *Nassella manicata** | Perennial |
| *Stipa megapotamia* | *Jarava filifolia** or *Nassella megapotamia*° | Perennial |
| *Stipa melanosperma* | *Nassella melanosperma** | Perennial |
| *Stipa mucronata* | *Nassella mucronate** | Perennial |
| *Stipa nardoides* | *Nassella nardoides** | Perennial |
| *Stipa neesiana* | *Nassella neesiana** | Perennial |
| *Stipa pampeana* | *Nassella pampeana** | Perennial |
| *Stipa pfisteri* | *Nassella pfisteri** | Perennial |
| *Stipa pulchra* | *Nassella pulchra** | Perennial |
| *Stipa rosengurttii* | *Nassella rosengurttii** | Perennial |
| *Stipa rupestris* | *Nassella rupestris** | Perennial |
| *Stipa sanluisensis* | *Nassella sanluisensis** | Perennial |
| *Stipa sellowiana* | *Nassella sellowiana** | Perennial |
| *Stipa tenuis* | *Nassella tenuis* if *Stipa tenuis* (Phil.).* or<br>*Aristida ternipes* if *Stipa tenuis* Willd. ex Steud.*<br>Both support perennial. | Perennial |
| *Stipa tenuissima* | *Nassella tenuissima** | Perennial |
| *Stipa tucumana* | *Nassella tucumana** | Perennial |
| *Stipa viridula* | *Nassella viridula** | Perennial |
| *Streblochaete longiarista* | *Koordersiochloa longiarista** | Perennial* |
| *Sylvipoa queenslandica* | Accepted name* | Perennial*, collected under *Poa queenslandica* |
| *Thamnocalamus tessellatus* | *Bergbambos tessellate** | Perennial |
| *Thinopyrum caespitosum* | *Elymus nodosus subsp. caespitosus** | Perennial (genus,*°) |
| *Thinopyrum intermedium* subsp. *intermedium* | Accepted name* | Perennial*, collected under *Elymus hispidus* |
| *Triarrhena lutarioriparia* | *Miscanthus lutarioriparius** | Perennial |
| *Trikeraia hookeri* | Accepted name* | Perennial*, collected under *Stipa hookeri* |
| *Trikeraia pappiformis* | Accepted name* | Perennial*, collected under *Stipa pappiformis* |
| *Yushania alpina* | *Oldeania alpina** | Perennial |
| *Zeugites americana* | *Zeugites americanus* | Perennial |
| *Zeugites latifolia* | *Zeugites latifolius* | Perennial |
| *Zeugites munroana* | *Zeugites munroanus* | Annual |
| *Zeugites sagittata* | *Zeugites sagittatus* | Perennial |
| *Zeugites smilacifolia* | *Zeugites smilacifolius* | Perennial |

**Table S2.** Log-likelihoods and AICc values of HMM models [2] with different numbers of rate categories.

| **Hidden rates model** | **Log likelihood** | **AICc** | **$\Delta$AICc** |
|---|---|---|---|
| **One-rate** | -1251.68 | 2507.36 | 170.08 |
| **Two-rate** | -1179.75 | 2371.53 | 34.25 |
| **Three-rate** | -1156.60 | 2337.28 | 0 |
| **Four-rate** | -1149.93 | 2340.10 | 2.82 |
| **Five-rate** | -1143.39 | 2347.32 | 10.04 |

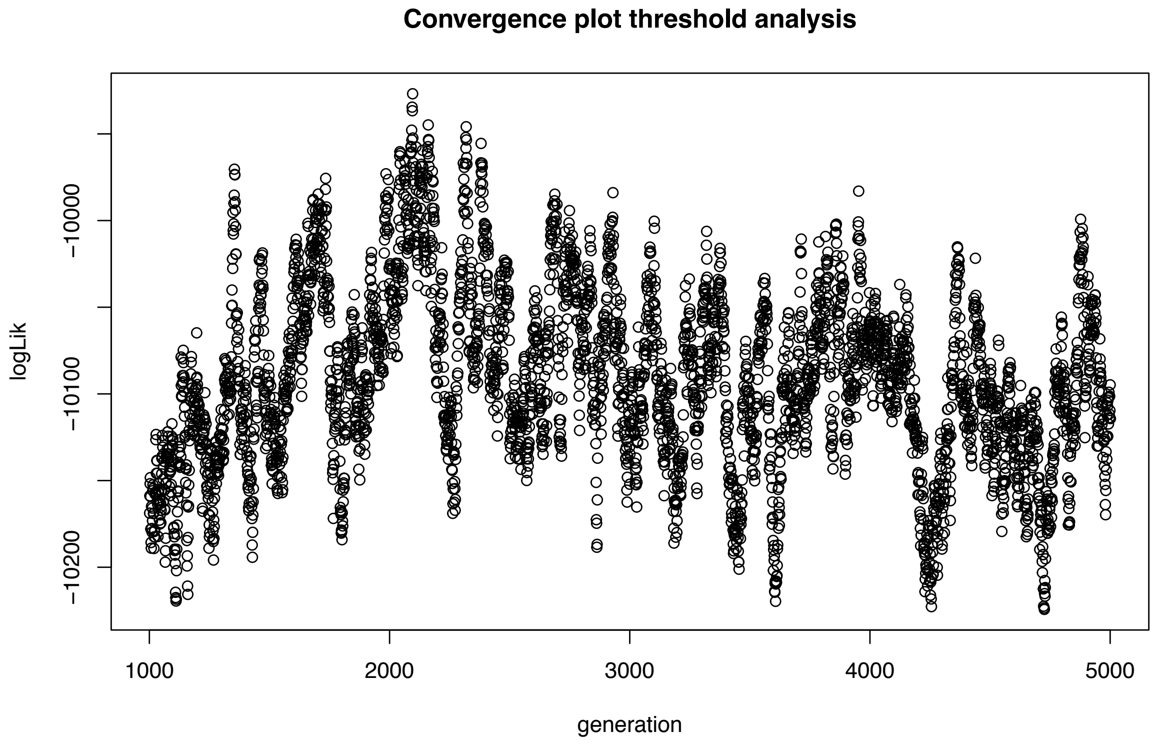


Figure S1. Convergence plot for the ancestral state reconstruction using the threshold model [3], showing log-likelihood (logLik) values across 5 million generations. The initial 20% of generations were discarded as burn-in. The liability factor was modelled as evolving under Brownian motion (BM). Other models (Ornstein-Uhlenbeck and Lambda) were tested as well, but these failed to converge after 5,000,000 generations, suggesting they are too complex for a dataset of this size. Most implementations of this method in the literature have also modelled the liability factor under BM.

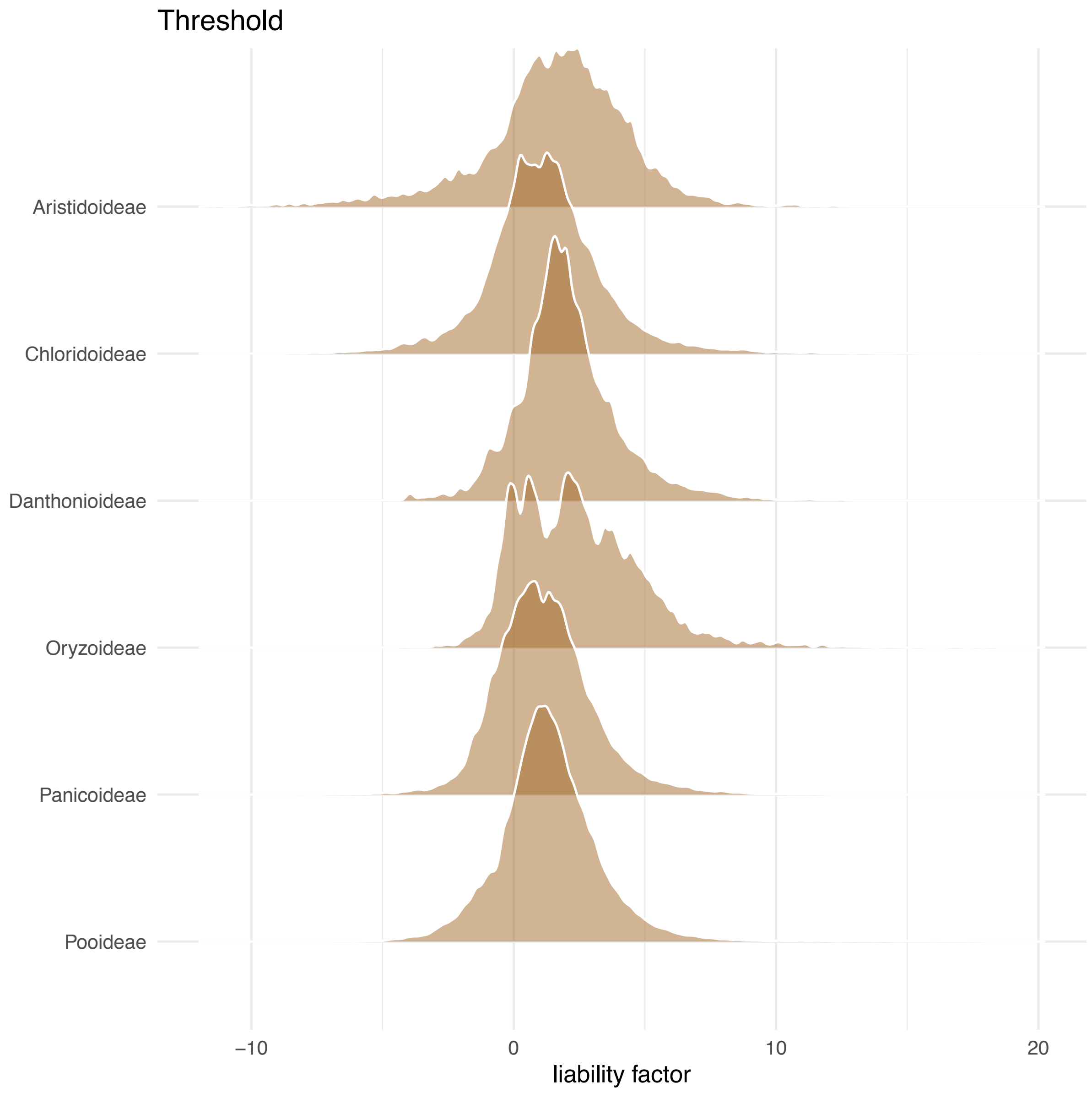


Figure S2. Smoothed histograms of liability factors at the nodes sampled during the ancestral state reconstruction using the threshold model [3], or the six sampled subfamilies. Nodes with values closer to the threshold (0) are more liable to change between the annual (negative values) and perennial (positive values) states.

### Text S1. Supplementary Discussion

#### Ancestral state reconstructions under the hidden rates and threshold models

The two ancestral state reconstructions (hidden rates model and threshold model) showed widely different patterns: many origins (n=63) and few reversals (n=13) under the hidden rates model, compared to the reverse pattern under the threshold model (19 vs. 32). These differences can partly be explained by higher levels of certainty under the hidden rates model (except in the Panicoideae, **figures 1, 2**, main article). Some differences may also be due to the different number of rate categories used in these models (3 vs. 1). But which is a more accurate representation of the evolutionary history of life history in grasses? Considering the size of our dataset, likely necessitating incorporation of rate heterogeneity [4], and the unrealistic estimates of origins and reversals under the threshold model, we consider the hidden rates model more reliable. This contradicts a previous comparison between these two methods [5], albeit on a different and much smaller dataset.

#### Life history evolution is stable in Pooideae

The finding of trait stability in Pooideae is particularly interesting, given the numerous independent origins of annuals, concentrated in a core clade with different resource allocation patterns compared to the rest of the clade [6]. This type of pattern may easily have been interpreted as evidence for life history being inherently labile in this clade. However, when tested against expectations under stable and labile scenarios (**table 2, figure 3**, main article), or in the context of a broader clade (**figure 1,** main article), these patterns were clearly inferred as stable. Inference of evolutionary stability for this clade is not at odds with a high number of (independent) origins of annuals – Pooideae is the largest subfamily of grasses, comprising >4100 species [7]. The second largest clade, Panicoideae (>3300 species), showed a very different distribution of annuals and life history evolution was inferred to be labile in this clade. Perhaps *because* most life history shifts in Pooideae are concentrated in the core clade, we infer trait stability for Pooideae as a whole, while a separate test *within* core Pooideae (the 'precursor clade' of Lindeberg et al. [6]) could still reveal lability in this clade specifically.
